# Snake venom metalloproteinases are predominantly responsible for the cytotoxic effects of certain African viper venoms

**DOI:** 10.1101/2024.12.04.626778

**Authors:** Keirah E. Bartlett, Adam Westhorpe, Mark C. Wilkinson, Nicholas R. Casewell

## Abstract

Snakebite envenoming is a neglected tropical disease that causes substantial mortality and morbidity globally. The puff adder (*Bitis arietans*) and saw-scaled viper (*Echis romani*) have cytotoxic venoms that cause permanent injury via tissue-destructive dermonecrosis around the bite site. Identification of cytotoxic toxins within these venoms will allow development of targeted treatments, such as small molecule inhibitors or monoclonal antibodies to prevent snakebite morbidity. Venoms from both species were fractionated using gel filtration chromatography, and a combination of cell-based cytotoxicity approaches, SDS-PAGE gel electrophoresis, and enzymatic assays were applied to identify venom cytotoxins in the resulting fractions. Our results indicated that snake venom metalloproteinase (SVMP) toxins are predominately responsible for causing cytotoxic effects across both venoms, but that the PII subclass of SVMPs are likely the main driver of cytotoxicity following envenoming by *B. arietans*, whilst the structurally distinct PIII subclass of SVMPs are responsible for conveying this effect in *E. romani* venom. Identification of distinct SVMPs as the primary cytotoxicity-causing toxins in these two African viper venoms will facilitate the future design and development of novel therapeutics targeting these medically important venoms, which in turn could help to mitigate the severe life and limb threatening consequences of tropical snakebite.

**Key Contribution:** SVMP toxins were identified as the primary cytotoxicity-causing toxins in the venoms of the puff adder (*Bitis arietans*) and saw-scaled viper (*Echis romani*); PII and PIII SVMPs, respectively. This cytotoxicity can be prevented using the metalloproteinase-inhibiting chelator EDTA, suggesting targeted drugs/antibodies may be a viable option for future treatment.

## 1. Introduction

Snakebite envenoming is a neglected tropical disease (NTD) with a high mortality and morbidity burden in rural regions of sub-Saharan Africa, South and South-east Asia, and Latin America [1]. Causing up to 138,000 deaths and 400,000 disabilities annually, snakebite has been designated a World Health Organization (WHO) priority NTD since 2017, and an ambitious target has been set aiming to halve the number of deaths and disabilities attributed to snakebite by 2030 [1,2]. Currently, the only specific therapeutic for snakebite is antivenom and whilst invaluable in saving lives, this treatment often carries with it a range of affordability, efficacy, and safety issues [1,3]. Consequently, several new therapeutics are under investigation, such as monoclonal antibodies and small molecule inhibitors [4–7], for potently inhibiting key toxins within medically important venoms. However, it is critical that we first understand which venom components are of greatest pathogenic importance in medically important venoms to guide the discovery and development of more targeted and efficacious treatments [8].

Whilst some venoms cause life threatening systemic effects, such as haemorrhage or respiratory paralysis, many patients experience severe local tissue damage in the region surrounding the bite, which often results in permanent sequelae such as limb or digit impairment and/or the requirement for surgical interventions such as debridement of necrotic tissue [9,10]. In Africa alone, as many as 14,600 snakebite patients may require limb amputation in response to snakebite-induced local tissue damage each year [2,11]. In Africa, localised tissue necrosis following snakebite can largely be attributed to venoms from a range of spitting cobras (family Elapidae) and vipers (family Viperidae) [12]. The vipers primarily responsible for causing morbidity following envenoming are the puff adder (*Bitis arietans*) and the saw-scaled vipers (e.g. *Echis ocellatus*, *E. romani*, *E. pyramidum* and others) [12,13]. In West Africa, *E. romani* (recently reclassified and previously known as *E. ocellatus* [14,15]) is one of the main biting species, and in Nigeria alone it contributes to approximately 90% of severe envenomings [16]. Collectively, *Bitis arietans* are believed to cause a substantial number of severe envenomings in eastern, central, and southern Africa [12]. Both these venoms are characterised by causing extensive local tissue damage as the result of venom cytotoxicity, in addition to inducing systemic haemostatic disturbances such as haemorrhage, coagulopathy, and hypotension [13,17].

Like many snake venoms, those of *E. romani* and *B. arietans* are complex, consisting of a diverse mixture of toxin isoforms from several distinct protein families. Previous transcriptomic and proteomic characterisations show that these venoms consist predominantly of snake venom metalloproteinases (SVMPs), phospholipases A2 (PLA2), C-type lectin-like proteins (CLPs), disintegrins, snake venom serine proteases (SVSPs), cysteine-rich secretory proteins, L-amino acid oxidases (LAAOs), and kunitz-type serine protease inhibitors [18–22]. However, the venom compositions of these species vary considerably. For example, in Nigeria the most abundant venom proteins in *B. arietans* venom are CLPs, followed by SVMPs [22], while Nigerian *E. romani* venom is composed mostly of SVMPs, with CLPs, PLA2 and serine proteases at lower levels [19,21].

Whilst both *E. romani* and *B. arietans* venoms cause cytotoxicity, the toxins responsible for causing this morbidity remain unclear. Previous studies have shown that members of the SVMP, PLA2 and LAAO snake venom toxin families can all exhibit cytotoxic effects [1,23,24]. In addition to their high relative abundance in many viper venoms, SVMPs (which are zinc-dependent) are associated with causing diverse functional activities presenting as local and systemic haemorrhage, coagulopathy, oedema, myonecrosis, blistering, and dermonecrosis in snakebite patients [25–27]. The SVMPs are structurally classified into three main groups, the PI, PII, and PIII sub-classes, with PI SVMPs containing only a metalloproteinase domain, the PIIs a metalloproteinase and a disintegrin domain, and the PIIIs a metalloproteinase, a disintegrin-like, and a cysteine-rich domain [28]. Each of these subclasses cause distinct pathology in snakebite patients, with PIs considered to be mostly fibrinogenolytic, PIIs predominantly inhibiting platelet aggregation, and PIIIs causing haemorrhage and coagulopathy [23]; however, a previous study by Freitas-de-Sousa *et al.* [29] showed dermonecrosis following envenoming by the South American pit viper *Bothrops atrox* was attributed to both PI and PIII SVMPs, which were among the most abundant toxins found in the venom [30]. While there are multiple groups of PLA2, only two of these classes are found in snakes: class I in elapids and colubrids, and class II in vipers [31]. Viperid PLA2 are known to cause local and systemic myotoxicity, cytotoxicity, and necrosis of skeletal muscle, with numerous studies reporting this from *Bothrops* species [9,32–37]. L-amino acid oxidases are known to contribute towards venom toxicity and can be cytotoxic and apoptotic, although there is not yet a clear consensus on the exact mechanisms responsible for this [38]. Costal-Oliveira *et al.* [39] showed LAAO from *B. atrox* caused cytotoxicity *in vitro* using human epidermal keratinocytes, while LAAO from *Cerastes cerastes* venom has been shown to cause necrosis *in vivo* [40].

As a first step towards target identification of morbidity-causing toxins found in medically important African vipers, in this study we sought to identify the toxins responsible for causing venom-induced cytotoxicity induced by *B. arietans* and *E. romani*. To do so, we employed a combination of cell and enzyme based *in vitro* assays coupled with chromatographic profiling and separation of the crude snake venoms into their toxin constituents. Our findings demonstrate that SVMP toxins are largely responsible for the cytotoxic effects of both snake venoms observed in cellular assays using human epidermal keratinocytes; however, we find that PIII isoforms are largely responsible for the cytotoxic effects of venom from the saw-scaled viper *E. romani*, while PII SVMPs underpin venom cytotoxicity caused by the puff adder *B. arietans*.

## 2. Results

### 2.1 Saw-scaled viper and puff adder venoms cause cell cytotoxicity in human epidermal keratinocytes

We first investigated the cytotoxic potential of *E. romani* and *B. arietans* venoms in a previously described cell cytotoxicity assay using immortalised human epidermal keratinocytes (HaCaT cell line) [41,42]. The HaCaT cells were exposed to crude venoms at various concentrations for 24 hours followed by the addition of thiazole blue tetrazolium (MTT) assaying to assess cell viability via measures of metabolic activity [43,44]. Both venoms were potently cytotoxic *in vitro*, although IC50 analyses revealed that *E. romani* venom was slightly more cytotoxic than the venom of *B. arietans* (IC50 7.8 µg/mL ± 2.7 vs IC50 17.8 µg/mL ± 6.2, respectively) (**Fig. 1**).

**Fig. 1.**
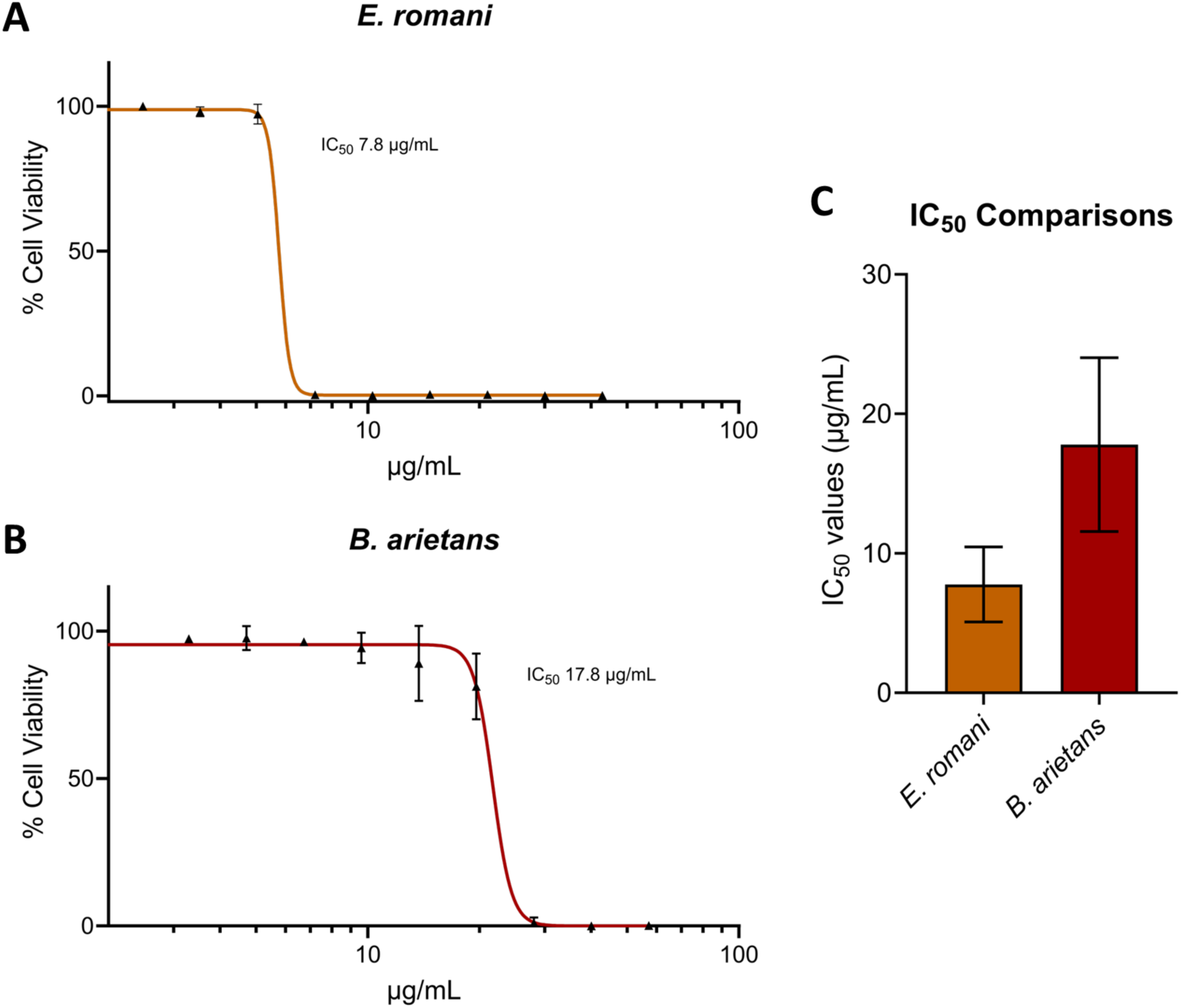
Crude venoms from the saw-scaled viper and puff adder affect cell viability. Cell viability was measured in immortalised human keratinocytes (HaCaT cells) using MTT assays. HaCaT cells were treated for 24 h with serial dilutions of *E. romani* (**a**) and *B. arietans* (**b**) venoms, displayed here as 95% confidence bands. (**c**) IC50 summary of venom concentration curves displayed in a and b. For panels a and b, the data shown represent mean % cell viability and corresponding standard deviations. All data displayed are from three independent experiments with each condition conducted in triplicate. Data were normalised to 0-100% between the lowest and highest read values for analysis, then plotted as concentration-response curves using GraphPad Prism 9. For panel c, the data shown represents the mean IC50 values of curves in a-b and corresponding standard deviations.

### 2.2 Venom fractions responsible for causing cell cytotoxicity are rich in SVMP toxins

To investigate which toxins in *E. romani* and *B. arietans* venoms are responsible for causing epidermal cell cytotoxicity, we used a chromatographic approach to separate crude venoms into fractions before retesting these fractions for the effects against the HaCaT cell line via MTT assays. Venoms were separated into fractions by carrying out size exclusion chromatography (SEC) under native conditions. The resulting MTT assays showed that *E. romani* venom contained several adjacent venom fractions (fractions 14-19) that exhibited cytotoxic effects, with significant reductions in cell viability detected to between 0.6% (± 0.9) and 8.8% (± 14.5) (*P*=<0.0001) of the vehicle control (**Fig. 2a**). When venom fraction cytotoxic activities were aligned with the chromatographic traces generated from the fractionation process, the active fractions in the MTT assay appeared to correlate with the abundant amounts of PIII SVMPs found in this venom (**Fig. S1**). To confirm this hypothesis, each fraction was subsequently run on SDS-PAGE to determine the molecular mass of the toxins present (**Fig. 2b**). The cytotoxic-active fractions further correlated with the presence of PIII SVMPs, as each was shown to contain multiple protein bands ranging from ∼50-80 kDa in size. This analysis also revealed the interesting finding that the *E. romani* PI SVMP visible in fractions 25 and 26 on the gel (∼25 kDa) did not exert cytotoxic effects on epidermal keratinocytes.

**Fig. 2.**
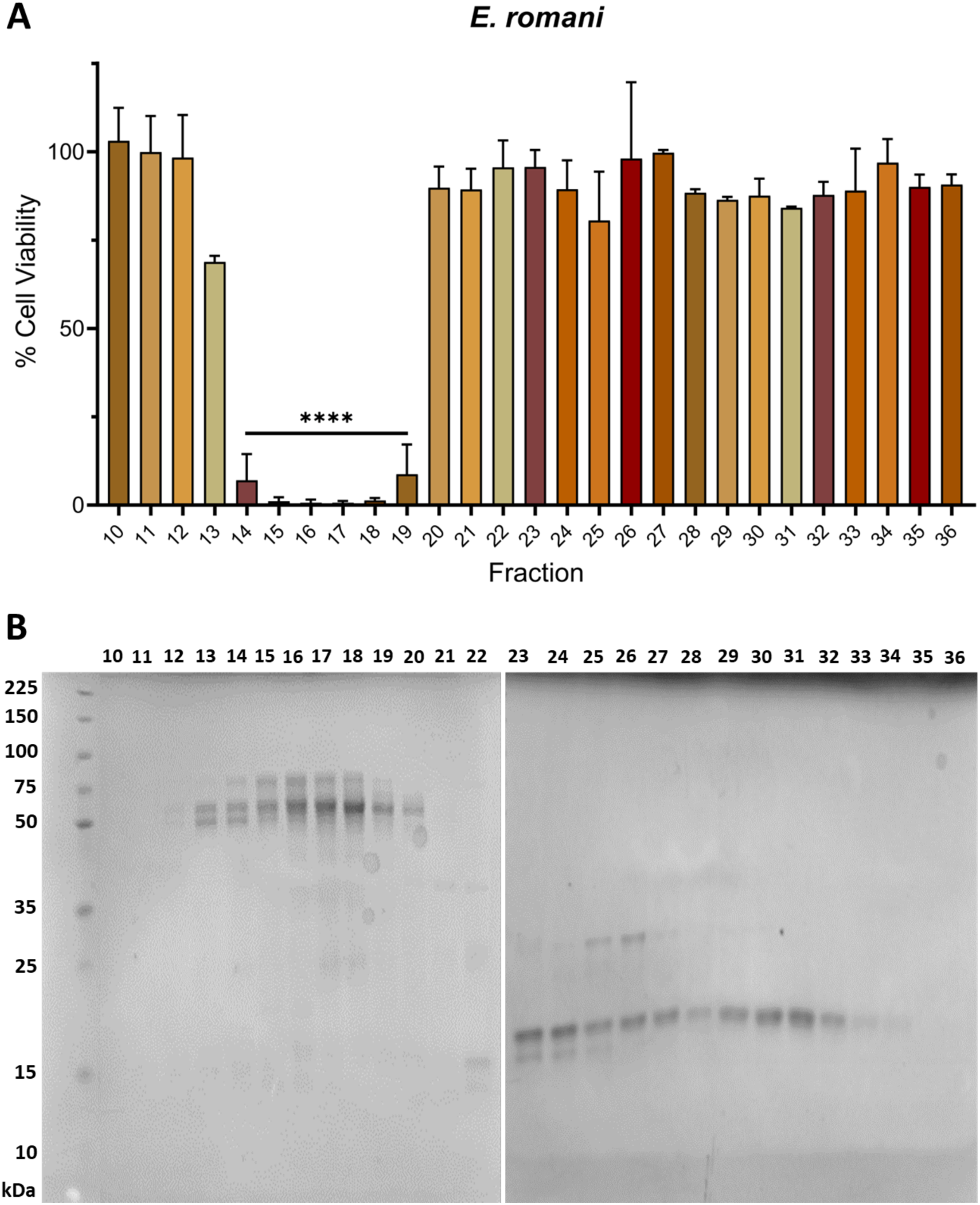
Cytotoxic venom fractions from *E. romani* correlate with the presence of PIII SVMP toxins. **a**) Cell viability was measured in immortalised human keratinocytes (HaCaT cells) using MTT assays. HaCaT cells were treated with fractions of Nigerian *E. romani* venom, and **b**) visualised using reducing SDS-PAGE gel electrophoresis on two separate gels. For panel a, the data shown represent mean % cell viability and corresponding standard deviations. All data displayed are from three independent experiments with each condition conducted in triplicate. Data were plotted using GraphPad Prism 9. Statistically significant differences were determined by one-way ANOVA, followed by a Dunnett’s multiple comparisons test, and are denoted by asterisks: **** (*P*<0.0001).

We used the same approach to identify which *B. arietans* SEC fractions cause cytotoxicity in the MTT assays. As with *E. romani*, multiple SEC fractions were responsible for causing cytotoxicity, with significant reductions in cell viability observed in two main clusters. Cell viabilities were reduced to 0.0% (± 0.9) and 0.1% (± 1.0) (*P*=<0.0001) for fractions 20- 22 and to between 0.0% (± 1.1) and 0.6% (± 0.4) (*P*=<0.0001) for fractions 27-29 (**Fig. 3a**). When aligned to SDS-PAGE analysis of the fractions, we found protein bands that corresponded with the molecular mass of the recently reported 34 kDa PIIa SVMP from Nigerian *B. arietans* venom (see fractions 20-23) and the 21 kDa IIa SVMP from Tanzanian *B. arietans* venom (see fractions 27-29) [45] (**Fig. 4.3B**). Additionally, there are multiple bands with mass of ∼15-20 kDa in fractions 23-32, which we speculate may include PLA2, disintegrins, or CLP monomers [21], that could potentially contribute to the observed cytotoxicity in fractions 27-29 **(Fig. 4.3B)**. Although there was no cytotoxic activity associated with fractions 18 and 19, which contains serine protease bands at 50-55 kDa [45], we did observe a modest, though still significant, reduction in cell viability in fraction 17 (51.0% ± 29.7; *P* = 0.022), which we hypothesise may contain LAAO, due to this fraction having a band at ∼60 kDa (**Fig. 4.3A** and **4.3B**).

**Fig. 3.**
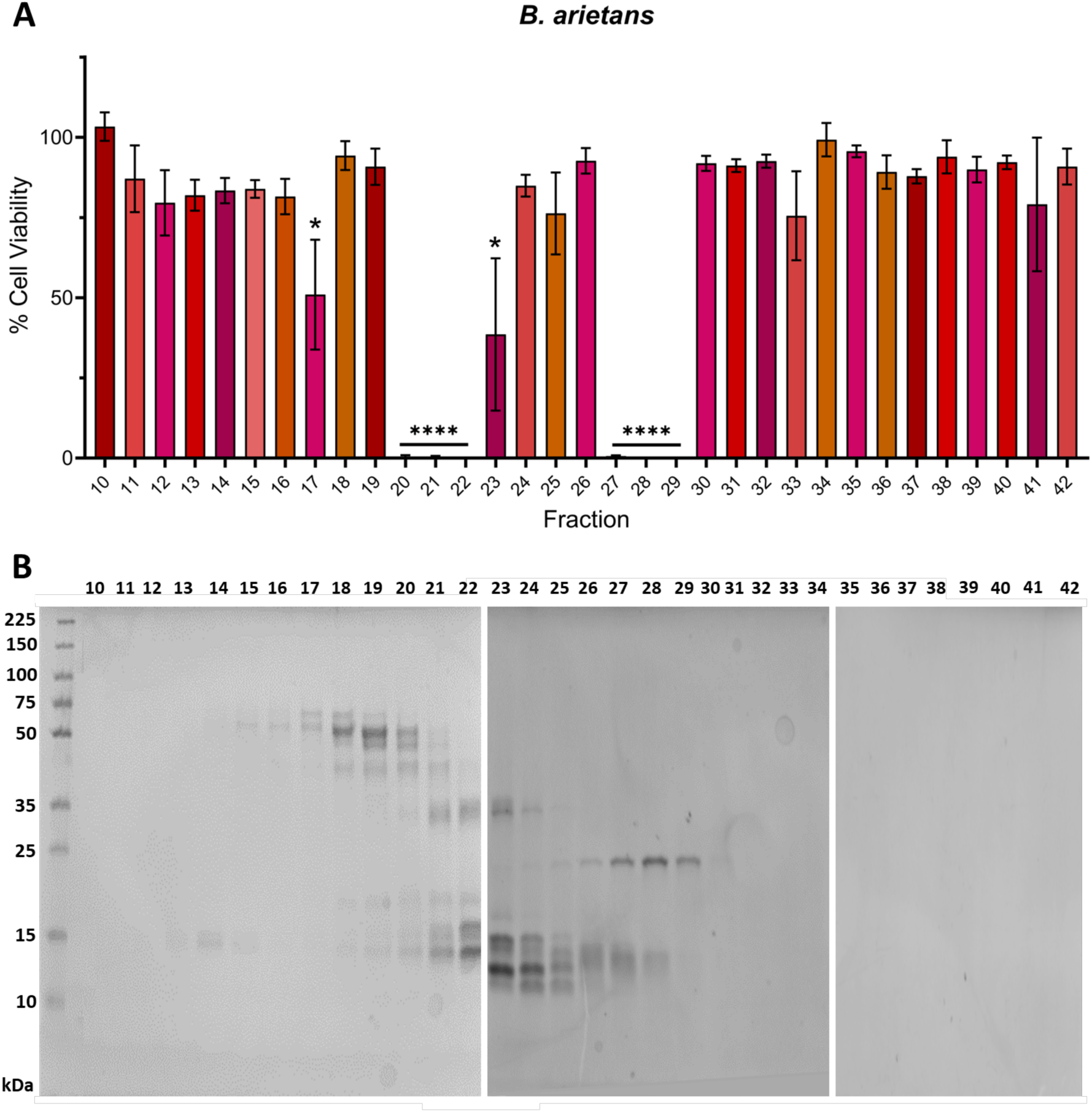
Cytotoxic venom fractions from *B. arietans* correlate with the presence of PII SVMP toxins. **a**) Cell viability was measured in immortalised human keratinocytes (HaCaT cells) using MTT assays. HaCaT cells were treated with fractions of *B. arietans* venom, and **b**) visualised using a reducing SDS-PAGE on three separate gels. For panel a, the data shown represent mean % cell viability and corresponding standard deviations. All data displayed are from three independent experiments with each condition conducted in triplicate. Data were plotted using GraphPad Prism 9. Statistically significant differences were determined by one-way ANOVA, followed by a Dunnett’s multiple comparisons test, and are denoted by asterisks: * (*P*<0.05), **** (*P*<0.0001).

**Fig. 4.**
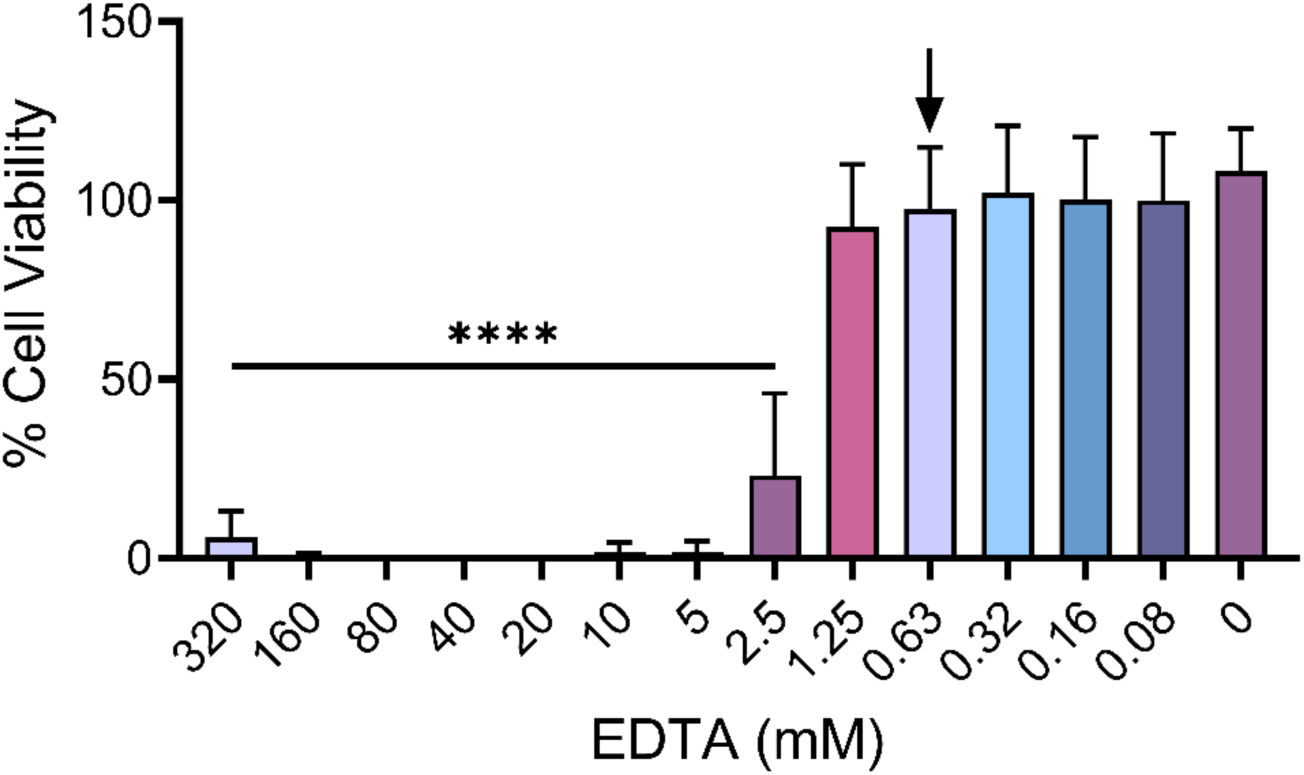
Establishing an appropriate concentration of EDTA to use in cell viability experiments. Cell viability was measured in immortalised human keratinocytes (HaCaT cells) using MTT assays. HaCaT cells were treated for 24 h with serial dilutions of EDTA. The maximal concentration select for use as treatment in subsequent inhibition assays was determined to be 0.625 mM EDTA (black arrow; concentrations displayed in axis are rounded to 2 dp). The data shown represent mean % cell viability and corresponding standard deviations. All data displayed are from three independent experiments with each condition conducted in triplicate. Data were plotted using GraphPad Prism 9. Statistically significant differences were determined by one-way ANOVA, followed by a Dunnett’s multiple comparisons test, and are denoted by asterisks: **** (*P*<0.0001).

### 2.3. The metalloproteinase inhibitor EDTA significantly reduces cytotoxic effects of venom fractions

To further explore the above findings, which suggested that SVMPs play a key role in *B. arietans* and *E. romani* cytotoxic effects, we next performed venom inhibition experiments with the metalloproteinase-inhibiting metal chelator EDTA, which chelates zinc and has previously been used in *in vitro* neutralisation tests and *in vivo* experiments investigating protection against venom-induced haemorrhage and dermonecrosis [46–49]. First, to understand how much EDTA could be used without affecting the viability of the HaCaT cell line, we conducted MTT assays and determined the highest EDTA concentration that could be used for downstream inhibition experiments. We prudently selected the second highest concentration that did not lead to significant reductions in cell viability, which was determined to be 0.625 mM (**Fig. 4**).

Next, we repeated the earlier MTT assays in the presence of 0.625 mM EDTA to confirm whether cytotoxic activity in the fractions of both venoms was predominately caused by SVMP toxins. Cytotoxicity caused by *E. romani* venom fractions was reduced from six active fractions (fractions 14-19) causing significant reductions in cell viability (*P*=<0.0001) to only one fraction (fraction 16; 18.0% ± 31.1; *P* = 0.040) in the presence of EDTA, with some modest, yet non-significant, cytotoxicity remaining in the adjacent fractions (fractions 15 and 17) (**Fig. 5a and b**). Fraction 16 is that with the highest combined content of the ∼50-80 kDa proteins, and thus remaining activity may simply be the result of insufficient EDTA to inhibit complete activity of this fraction (**Fig. 2b**). To further explore this relationship between SVMP toxins and venom cytotoxicity, we performed an *in vitro* fluorescent-based substrate cleavage SVMP enzymatic activity assay [6,50] with the venom fractions. SVMP activity was mostly detected in fractions 15-18 (13.3-28.9 % RFU ± 16.2-39.0), with trace activity observed in fractions 11-14 (1.2-7.8% RFU ± 0.6-12.7) (**Fig. 5c**). Collectively, these findings demonstrate that cytotoxic fractions from *E. romani* venom contain SVMP toxins and that the cytotoxic activity of these fractions can mostly be inhibited by the SVMP-inhibiting metal chelator EDTA.

**Fig. 5.**
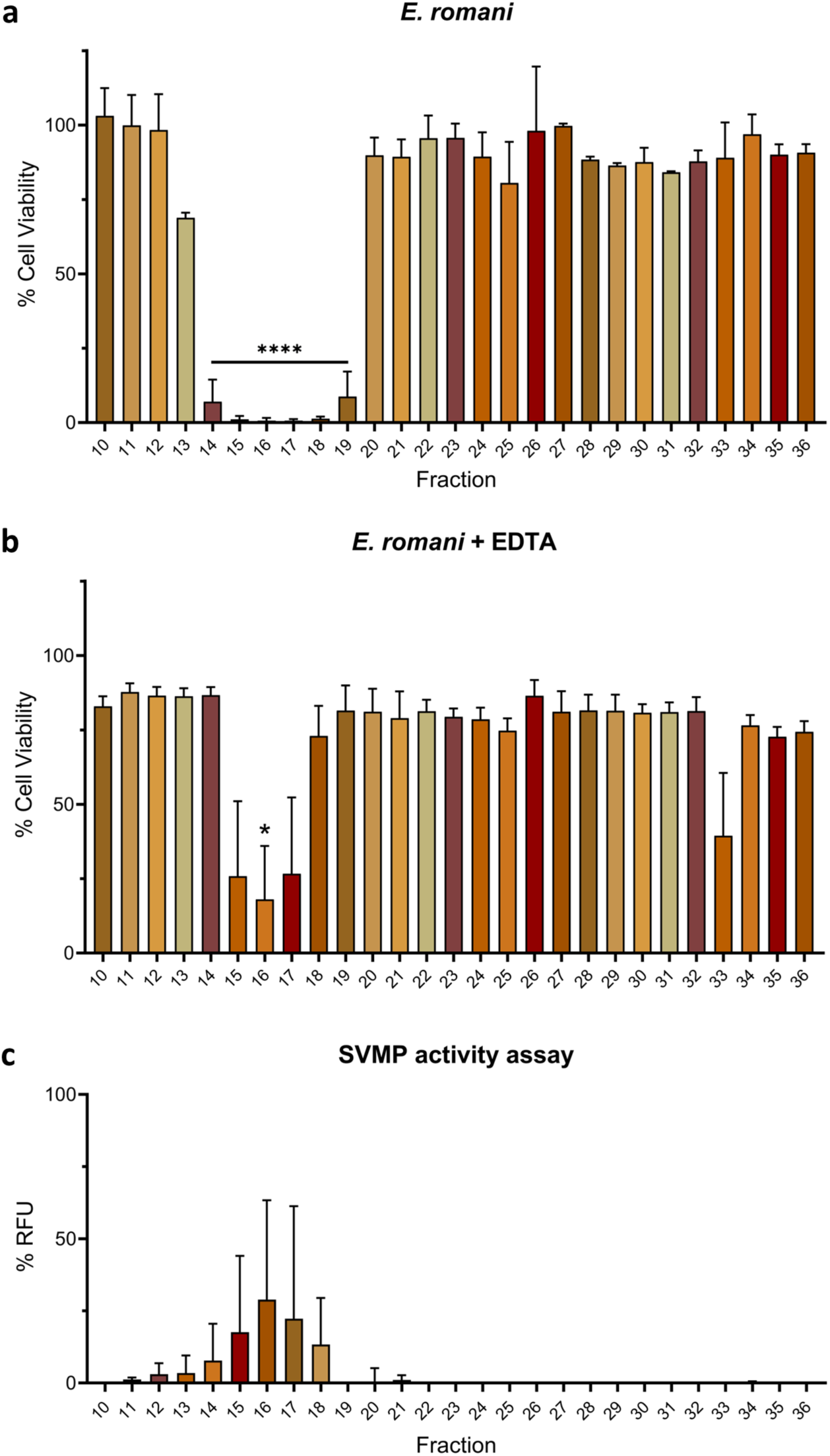
Cytotoxic fractions from *E. romani* venom exhibit SVMP activity and can be inhibited by the SVMP inhibiting metal chelator EDTA. Cell viability was measured in immortalised human keratinocytes (HaCaT cells) using MTT assays. HaCaT cells were treated with fractions of Nigerian **a**) *E. romani* venom, and **b**) *E. romani* venom in the presence of 0.625 mM EDTA. **c**) A fluorescent-based enzymatic SVMP activity assay showed that venom SVMP activity was observed in the same fractions that were active in the MTT assays (a) and were inhibited by EDTA (b). For panels a and b, the data shown represent mean % cell viability and corresponding standard deviations. The data displayed are from three independent experiments with each condition conducted in triplicate. For panel c, the data shown represent mean % RFU (intensity measured in relative fluorescence units), relative to crude venom (not shown), and corresponding standard deviations. The data displayed are from two independent experiments with each condition conducted in duplicate. Data were plotted using GraphPad Prism 9. For panels a and b, statistically significant differences were determined by one-way ANOVA, followed by a Dunnett’s multiple comparisons test, and are denoted by asterisks: * (*P*<0.05), **** (*P*<0.0001).

EDTA also abolished the activity of cytotoxic venom fractions from *B. arietans.* No reductions in cell viability were observed in the MTT assays for any venom fraction in the presence of 0.625 mM EDTA, which compared favourably to the initial assay where six venom fractions found in two clusters (fractions 20-22 and 27-29) resulted in potent cytotoxic effects (**Fig. 6a and b**). Similar to *E. romani*, findings from the *in vitro* SVMP activity assay correlated with the MTT assays, with multiple cytotoxic venom fractions displaying SVMP activity, most noticeably fractions 21-23 (6.1-10.4 % RFU ± 0.9-1.9) and fractions 27-29 (39.8-52.3 % RFU ± 1.2-12.4) (**Fig. 6c**). Low levels of SVMP activity were also observed in fractions 19, 20, 24-26, and 30 (all ≤5.6% RFU), all of which are adajcent to fractions previously identified as being cytotoxic, though these fraactions themselves did not cause significant reductions in cell viability as measured by the MTT assays (with the exception of fraction 20) (**Fig. 6**). These results stongly suggest that PII SVMPs largely underpin the cytotoxic effects of *B. arietans* venom observed in human keratinocytes. However, the identity and relative contribution of fraction 17 to venom cytotoxicity remains unclear; though only modestly cytotoxic, the activity of this fraction was effectively inhibited by EDTA, but did not display activity in the SVMP substrate assay (**Fig. 6**).

**Fig. 6.**
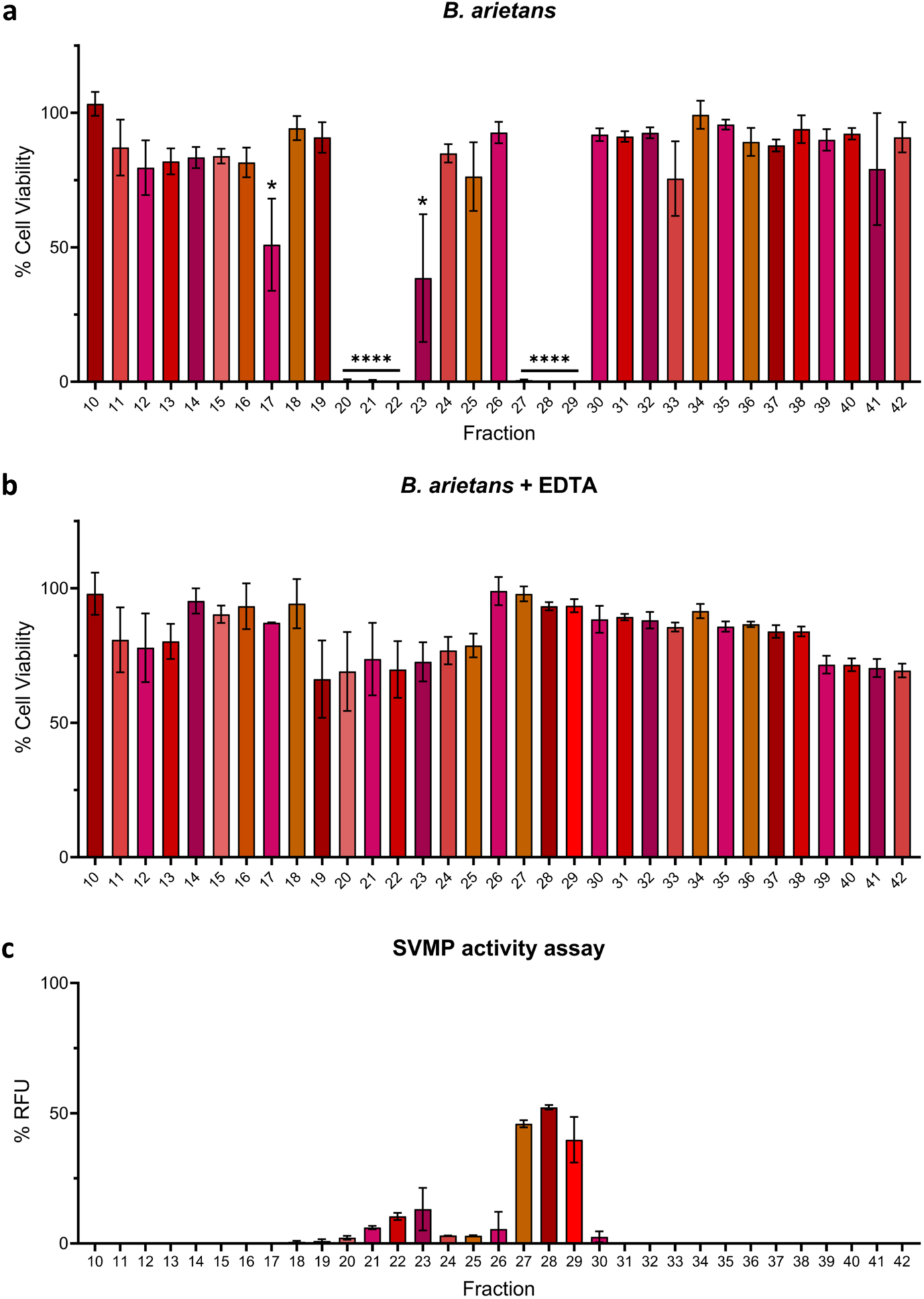
Cytotoxic fractions from *B. arietans* venom mostly exhibit SVMP activity and can be inhibited by the SVMP inhibiting metal chelator EDTA. Cell viability was measured in immortalised human keratinocytes (HaCaT cells) using MTT assays. HaCaT cells were treated with fractions of **a**) *B. arietans* venom, and **b**) *B. arietans* venom in the presence of 0.625 mM EDTA. **c**) A fluorescent-based enzymatic SVMP activity assay showed that venom SVMP activity was observed in most of the fractions that were active in the MTT assays. For panels a and b, the data shown represent mean % cell viability and corresponding standard deviations. The data displayed are from three independent experiments with each condition conducted in triplicate. For panel c, the data shown represent mean % RFU (intensity measured in relative fluorescence units), relative to crude venom (not shown), and corresponding standard deviations. The data displayed are from two independent experiments with each condition conducted in duplicate. Data were plotted using GraphPad Prism 9. For panel a, statistically significant differences were determined by one-way ANOVA, followed by a Dunnett’s multiple comparisons test, and are denoted by asterisks: **** (*P*<0.0001), * (*P*<0.05).

### 2.4. PIIa SVMPs from puff adders of differing geographical origin exhibit substantial differences in cell cytotoxicity

*Bitis arietans* venom is known to vary intraspecifically between different geographical regions [22,51]. Because we used a pool of venom from Nigerian and Tanzanian specimens for the experiments performed above, and in light of recent data demonstrating distinct substrate specificities of SVMPs sourced from East and West Africa *Bitis arietans* venoms, we next isolated the different cytotoxic PIIa SVMPs found in our venom pool by a previously established chromatography approach [45]. Toxin purification showed that the SVMP in fractions 20-22 was characteristic of the recently described Nigerian PIIa SVMP, while the fractions 27-29 contained the PIIa characteristic of Tanzanian venom (see [45]).

Given the prior description of distinct activities conferred by these two toxins ([45]), we next assessed and compared the relative toxicity of these two isolated proteins against HaCaT cells using MTT assay measures of cytotoxicity. The Tanzanian type PIIa was demonstrated to be significantly more cytotoxic than the Nigerian type PIIa, as demonstrated by resulting IC50 values (IC50 2.7 µg/mL ± 0.7 vs IC50 16.2 µg/mL ± 4.1; *P* = 0.005) (**Fig. 8**).

**Fig. 7.**
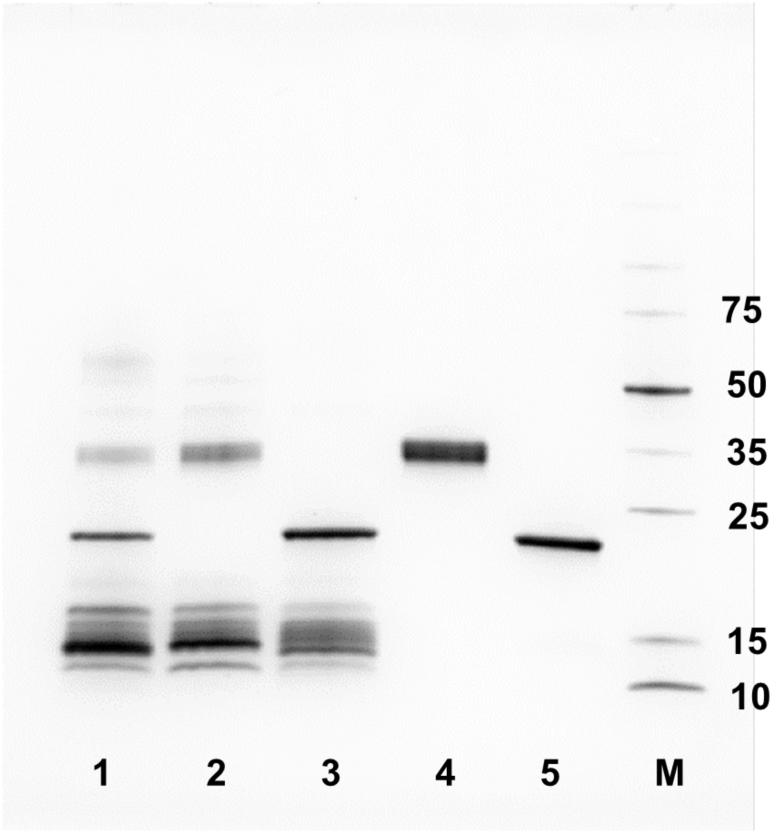
SDS-PAGE analysis of the *B. arietans* venom samples and purified SVMPs. The gel is 4-20% acrylamide (BioRad) and was run under reducing conditions. Eight μg of whole venom or 1-2 μg SVMP was loaded per lane and the gel was stained using Coomassie Blue R250. Lane 1, pooled venom; lane 2, Nigerian venom; lane 3, Tanzanian venom; lane 4, Nigerian PIIa SVMP (2 μg); lane 5, Tanzanian PIIa SVMP (1 μg); lane M, markers (Thermo Broad Range) with molecular weights indicated in kDa.

**Fig. 8.**
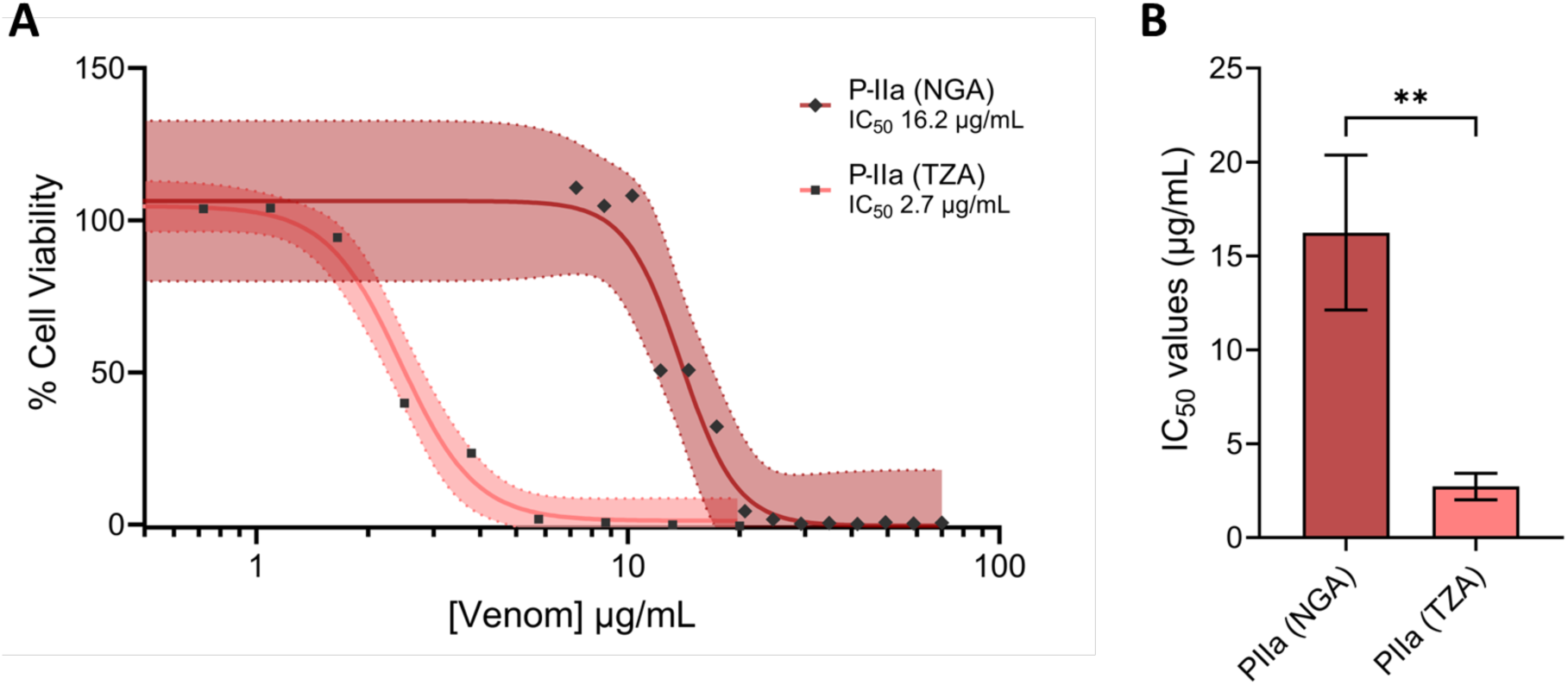
PIIa SVMPs from puff adder venom exhibit differences in cytotoxic potency. Cell viability was measured in immortalised human keratinocytes (HaCaT cells) using MTT assays. HaCaT cells were treated for 24 h with serial dilutions of PIIa SVMPs from Nigerian (NGA) and Tanzanian (TZA) *B. arietans* venoms, displayed here as **a**) 95% confidence bands, and **b**) IC50 values summary of venoms displayed in a. The data shown represent mean % cell viability and corresponding standard deviations. All data displayed are from three independent experiments with each condition conducted in triplicate. For panel a, data were normalised to 0-100% between the lowest and highest read values for analysis, then plotted as 95% confidence band curves using GraphPad Prism 9. For panel b, the data shown represents the mean IC50 values of curves in a and corresponding standard deviations. For panel b, statistically significant differences were determined by unpaired t-tests and are denoted by asterisks: ** (*P*<0.01).

### 2.5. The metalloproteinase inhibitor EDTA significantly reduces cytotoxic effects of whole venoms

As final confirmation that SVMPs are the primary cytotoxic components *in vitro*, the earlier MTT assays using whole crude venoms sourced from *E. romani* and *B. arietans* were repeated, but this time in the presence of EDTA. HaCaT cells were exposed to serial dilutions of crude venom in the presence of 0.625 mM EDTA, before cell viabilities were measured as described previously. Cytotoxicity was significantly reduced for both *E. romani* (IC50 7.8 µg/mL ± 2.7 vs IC50 16.5 µg/mL ± 0.5; *P* = 0.005) and *B. arietans* (IC50 17.8 µg/mL ± 6.2 vs IC50 30.5 µg/mL ± 0.2; *P* = 0.024) venoms when incubated with the metalloproteinase inhibiting metal chelator EDTA (**Fig. 9**).

**Fig. 9.**
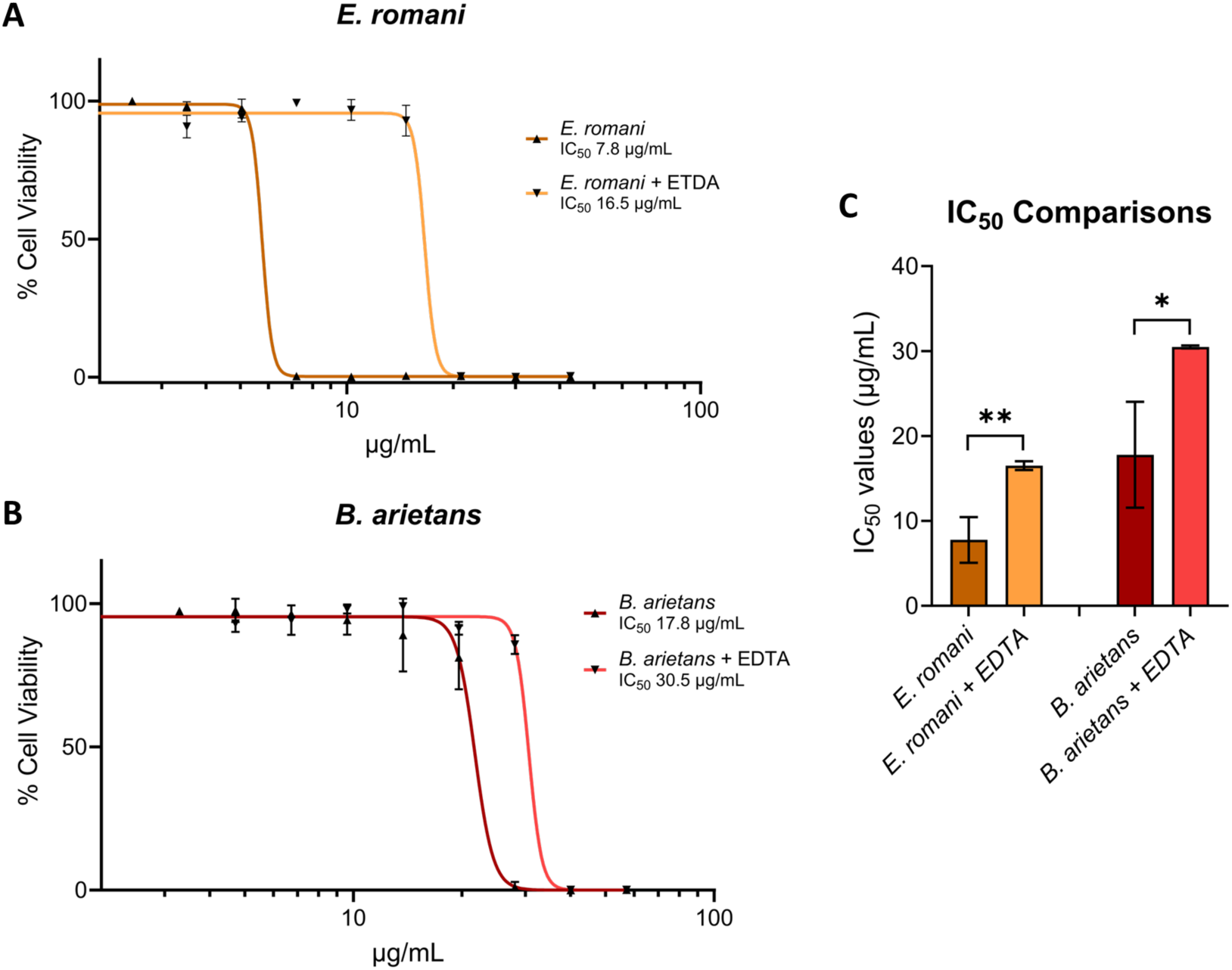
Cytotoxic crude venom activities are significantly reduced by the metalloproteinase inhibitor EDTA. Cell viability was measured in immortalised human keratinocytes (HaCaT cells) using MTT assays. HaCaT cells were treated for 24 h with serial dilutions of Nigerian **a**) *E. romani*, and **b**) *B. arietans* venoms, alone and with 0.625 mM EDTA, with the resulting data displayed as 95% confidence bands. **c**) IC50 values summary of venoms displayed in a and b using MTT assays. For panels a and b, the data shown represent mean % cell viability and corresponding standard deviations. All data displayed are from three independent experiments with each condition conducted in triplicate. Data were normalised to 0-100% between the lowest and highest read values for analysis, then plotted as concentration-response curves using GraphPad Prism 9. For panel c, the data shown represents the mean IC50 values of curves in a-b and corresponding standard deviations. For panel c, statistically significant differences were determined by unpaired t-tests and are denoted by asterisks: ** (*P*<0.01), * (*P*<0.05).

## 3. Discussion

Snakebite results in upwards of 2.5 million envenomings each year [52] and currently relies on antivenom as the sole therapeutic to treat patients [1]. Due to issues surrounding antivenom safety, efficacy, and affordability [3], work is ongoing to improve antivenom and develop alternative or adjunct treatments, such as monoclonal antibodies and small molecule inhibitors [53]. Such therapeutic modalities differ from mixtures of polyclonal antibodies by targeting specific toxin isoforms, and therefore it is critical to have a complete understanding of which venom toxins are of greatest pathogenic importance to target for neutralisation. Consequently, in this study we sought to understand which venom toxins present in saw-scaled viper and puff adder venoms are responsible for causing cytotoxicity, which is associated with contributing to severe local envenoming pathology and causing long term snakebite morbidity.

Using venoms from *E. romani* and *B. arietans*, we first used cell cytotoxicity assays with human epidermal keratinocytes to demonstrate that both venoms cause cytotoxic effects. This *in vitro* cytotoxicity has been documented previously and both venoms are known to be capable of causing local tissue necrosis in human patients [42,54,55]. To identify the toxins responsible for this medically relevant pathology, we used a chromatographic fractionation approach for both venoms, followed by cell cytotoxicity assays and SDS-PAGE gel electrophoresis. In both venoms, SVMPs were detected as the components present in the active fractions that were most likely to be cytotoxic, though different molecular masses indicated that PIII SVMPs are likely responsible for cytotoxic effects caused by *E. romani* venom, while the cytotoxic proteins detected in *B. arietans* venom were PII SVMPs. Inhibitory assaying further confirmed that SVMPs are the responsible toxins, as the metalloproteinase inhibitor EDTA significantly reduced the cytotoxic effects of the venom fractions and crude venom.

Previous studies using venom from *Bothrops* species have also shown that SVMPs are responsible for cytotoxicity and causing dermonecrosis. Freitas-de-Sousa *et al.* [29] found both the PI SVMP Atroxlysin-Ia and the PIII Batroxrhagin from *B. atrox* venom led to a rapid onset of dermonecrotic lesions in mice. Further, the small molecule SVMP inhibitors marimastat (a matrix metalloproteinase inhibitor) and DMPS (a metal chelator) have previously been shown to reduce *in vitro* cell cytotoxicity caused by *B. asper*, *Daboia russelii*, *E. carinatus*, *E. romani* and *Crotalus atrox* venoms, thereby hinting at broad SVMP-mediated toxicity [42]. Interestingly, no significant reduction in cytotoxicity was observed when *B. arietans* venom was treated with marimastat and DMPS [42], though this may simply reflect that a higher dose of these drugs is needed or that these inhibitors are less effective against PII toxins compared to PIII SVMPs. Additionally, the use of marimastat and DMPS *in vivo* have previously showed reductions in haemorrhagic or dermonecrotic dermal lesion sizes following intradermal injection of *C. atrox* or *E. ocellatus* (now reclassified as *E. romani*) venoms in mice [6], suggesting that, for these venoms at least, *in vitro* cytotoxicity experiments are potentially predictive of *in vivo* toxicity. Given our demonstration here that SVMPs are responsible for causing cytotoxicity caused by *B. arietans* venom, and that the SVMP-inhibiting metal chelator EDTA significantly reduces these cytotoxic effects, there may be considerable value in dose optimising ‘druggable’ inhibitors such as marimastat and DMPS [56] to rigorously explore their potential to prevent the local morbidity-causing effects of viper venoms. This is particularly pertinent given that different SVMP toxins were found to be responsible for toxicity in different puff adder samples, with the Tanzanian PIIa SVMP being significantly more cytotoxic than the PIIa found in Nigerian venom. While intraspecies venom variation has been previously reported in *Bitis arietans* [22], and also several other venomous snake species [57–62], the clinical implications of such variation remains unclear due to a scarcity of case reports describing puff adder envenomings in the literature [63]. It would be valuable to understand whether both these toxins cause severe local envenoming *in vivo*, and whether the differences in cytotoxic potency observed here *in vitro* result in distinctions in *in vivo* toxicity and/or the neutralising efficacy of toxin inhibitors or current antivenom therapy.

Although the EDTA significantly reduced the cytotoxic effects caused by *E. romani* and *B. arietans* crude venoms, some cytotoxicity remained. While it is possible that residual cytotoxicity may have been reduced further if higher amounts of EDTA were able to be used without causing cell toxicity, there is also the possibility that some components in the venom may work in a collective manner (i.e. additive/synergistic) to contribute to venom cytotoxicity, whether that be in combination with the detected cytotoxic SVMPs or other venom components, but that their cytotoxic effects may have been lost when the venom was fractionated into distinct constituents. We recently showed this for the venom of a spitting cobra (*Naja nigricollis*), where PLA2 toxins that were weakly cytotoxic by themselves enhanced the cytotoxic activity of other venom components when combined together [41]. It is possible that testing venom fractions for cytotoxicity may not completely represent the whole venom activity, and therefore recapitulating the fractionated venom minus the SVMP-containing fractions, or combining the bioactive fractions with other seemingly inactive fractions would be a robust follow-up study to further explore whether toxin potentiation occurs in *E. romani* and *B. arietans* venoms.

Despite the interesting findings presented in this study, there are several limitations. Firstly, we only investigated *in vitro* cytotoxicity. Our cell-based assay measures of cytotoxicity used a monolayer of keratinocytes, and therefore represents a simplistic model compared with in-development 3D organoid or organotypic models and *ex vivo* skin models, which may better recapitulate local envenoming *in vivo* via the combined complexity of multiple skin layers and cell types [41]. While SVMPs in both venoms were the predominant drivers of cytotoxicity in our cell-based assay, and therefore it is highly probable that these toxins would substantially contribute to causing dermonecrosis *in vivo*, other studies have highlighted distinctions between cell-based and *in vivo* findings, for the same reasons as outlined above [41]. Therefore, follow on preclinical experiments exploring the haemorrhagic and dermonecrotic activity of the isolated SVMP cytotoxins would be valuable to confirm these findings and to contextualise the therapeutic value of SVMP-inhibiting drugs. Finally, the work presented in this study focused on two viperid venoms from sub-Saharan Africa. Given that venom variation can be extensive across all taxonomic levels [64,65], and has previously been documented from different geographical locales of *B. arietans* [22,51], it would be valuable to perform similar experiments using venom sourced from specific locales across the broad geographical range of this species. Additionally, it might be informative to extend these studies to include other medically relevant *Echis* or *Bitis* species found throughout the continent.

In summary, our study has shown that SVMPs found in saw-scaled viper and puff adder venoms are largely responsible for causing venom cytotoxicity in cellular assays. While the toxins in these venoms are different, representing the PIII and PII structural subclasses of SVMP toxins in *E. romani* and *B. arietans* venoms, respectively, both sets of toxins are members of the same protein family and are inhibited by the metal chelator EDTA. The identification of key pathology-causing toxins, such as those described here, is essential for the rational design and development of novel therapeutics that seek to mitigate the severe life and limb threatening consequences of tropical snakebite.

## 5. Materials and Methods Materials

Thiazolyl blue methyltetrazolium bromide (MTT; M5655), NaHCO3 (401676), NaCl (S7653-1KG) and Coomassie brilliant blue G (B-0770) were purchased from Sigma (Sigma-Aldrich; Merck). Dulbecco’s modified Eagle’s medium (DMEM; 11574516), foetal bovine serum (FBS; 11573397), glutaMAX supplement (35050038), penicillin-streptomycin (11528876), phosphate buffered saline (PBS; 11503387), and TrypLE Express were purchased from Gibco (ThermoFisher Scientific). PageRuler Prestained Protein Ladder 10 to 180 kDa (26616), 2-mercaptoethanol (BP176), tween-20 (11417160), methanol (M/4000/PC17) and glycine (J64365.A1) were purchased from Thermoscientific (Fisher). EDTA (BIA4892) was purchased from Apollo Scientific Ltd. 4–20% Mini-PROTEAN TGX 10-well Precast Protein Gels (4561093) were purchased from BioRad. Gel loading dye (purple 6X, B7024S) was purchased from New England Biolabs. Tris base (30- 20-60) and SDS (30-33-60) were purchased from Severn Biotech Ltd. Acetic acid glacial (20104.334) was purchased from VWR. SVMP substrate (ES010) was purchased from Bio-Techne.

### Snake venoms

Venoms were sourced from snake specimens maintained in the Liverpool School of Tropical Medicine’s (LSTM) Herpetarium. This facility and its snake husbandry protocols are approved and inspected by the UK Home Office and the LSTM Animal Welfare and Ethical Review Boards. Venoms were extracted from wild-caught animals, namely: *Bitis arietans* (mixed batch containing venom from both Nigerian and Tanzanian specimens, 13 individuals), and *Echis romani* (Nigeria, 38 individuals in venom pool). Crude venoms were lyophilised and stored at 4 °C to ensure long-term stability. Venoms were resuspended at the desired concentrations in PBS (pH 7.4) prior to work.

### Venom toxin fractionation and isolation

The protein components of the venoms used in this study were separated using size exclusion chromatography (SEC). For this, 5 mg of venom (*Echis romani* or *Bitis arietans*) was dissolved in 0.5 mL ice-cold PBS (25 mM sodium phosphate pH 7.2, 0.15 M NaCl) and centrifuged at 10,000 xg for 10 minutes. The supernatant was immediately loaded onto a 24 mL column of Superdex 200HR (Cytiva) equilibrated in PBS. The column was operated at a flow rate of 0.2 mL/min during loading and 0.5 mL/min for elution. 0.5 mL fractions were collected after the void volume and elution was monitored at 214 and 280 nm. SDS-PAGE analysis was carried out on all fractions seen to contain protein on the trace. The fractions were stored at -20°C until use.

### Cell culture

The immortalised human epidermal keratinocyte line, HaCaT [66,67], was purchased from Caltag Medsystems (Buckingham, UK). Cells were cultured in phenol red-containing DMEM with GlutaMAX supplemented with 10% FBS, 100 IU/mL penicillin, 250 µg/mL streptomycin, and 2 mM sodium pyruvate (standard medium; Gibco), per Caltag’s HaCaT protocol. The cells were split and growth medium changed twice weekly up to a maximum of 30 passages. Cells were maintained in a humidified, 95% air/5% CO2 atmosphere at 37 °C (standard conditions).

### MTT measures of cell metabolic activity

To determine the cytotoxic effects of crude snake venoms and venom toxin fractions, MTT assays were performed. MTT cell viability [43] assays were completed as described previously [42], with minor modifications. Briefly, HaCaT cells were seeded at 20,000 cells/well in clear-sided/clear-bottomed 96-well plates in standard medium and incubated overnight in standard conditions. The following day, standard medium was aspirated and replaced with treatment solutions prepared in 1% FBS-DMEM medium. Wells were treated in triplicate with 100 µL of crude venoms sourced from *E. romani* (2.47-42.85 µg/mL) or *B. arietans* (3.29-57.14 µg/mL), then placed back in standard conditions for a further 24 hours. The treatment solutions were then aspirated and replaced with MTT-containing 1% FBS-DMEM medium (120 µL at 0.83 mg/mL), and the plates incubated for 45 min in standard conditions. Thereafter, the MTT-containing medium was aspirated, 100 µL DMSO added to each well to dissolve the formazan crystals, and absorbance (550 nm) was read for all wells on a CLARIOstar plate reader (BMG Labtech). Experiments were repeated independently on three occasions for each venom. Subsequently, data were normalised to 0-100% between the lowest and highest absorbance values for analysis to represent %-cell viability (MTT), then plotted as either concentration-response curves or as 95% confidence band curves using GraphPad Prism 9. IC50 values were calculated using the ‘[Inhibitor] vs. normalized response -- Variable slope’ -- Variable slope’ function.

The above process was repeated using venom fractions from each venom to identify cytotoxins, and then with isolated Nigerian and Tanzanian PIIa SVMP to compare the relative cytotoxicity between these two variants. Cells were seeded as described above and treated in triplicate with 100 µL of 1% FBS-DMEM medium containing neat fraction, at a 9:1 ratio of medium:fraction. Subsequent incubation steps, MTT treatments and measurements were carried out as described above. Experiments were repeated independently on three occasions for each fraction, with %-cell viability (non-normalised) data plotted as bar graphs using GraphPad Prism 9. Subsequently, purified Nigerian PIIa (7.26-70.00 µg/mL) or Tanzanian PIIa (0.72-20.00 µg/mL) SVMPs were tested in the format described above for crude venoms. Data were normalised to 0-100% between the lowest and highest absorbance values for analysis to represent %-cell viability and plotted as 95% confidence band curves, using the GraphPad Prism 9 function ‘[Inhibitor] vs. response -- Variable slope (four parameters)’.

### Inhibition of cytotoxic effects by EDTA

To assess whether venom SVMPs were responsible for cell cytotoxicity, we performed neutralisation experiments with the metalloproteinase-inhibiting metal chelator EDTA. First, to determine how much EDTA was tolerated by the cell line, HaCaT cells were seeded at 20,000 cells/well in 96-well plates in standard medium, as described above, and incubated overnight under standard conditions. Wells were treated in triplicate with 100 µL of 1% FBS-DMEM with varying concentrations of EDTA (78.1 µM – 320.0 mM). After 24 h incubation under standard conditions, MTT treatments and measurements were carried out as described above. Experiments were repeated independently on three occasions. Subsequently, data (non-normalised) were plotted as a bar chart using GraphPad Prism 9. From this, the maximal concentration of EDTA used in downstream assays was determined to be 0.625 mM, defined as the second highest concentration that did not significantly reduce cell viability.

Next, we tested whether EDTA could inhibit the SVMP action of both venom fractions and crude venoms. The previously described MTT assays were repeated with 0.625 mM EDTA added to each treatment, with MTT additions and %-cell viability readings carried out as above. Experiments were repeated independently on three occasions. Data for each fraction (non-normalised) were plotted as bar graphs using GraphPad Prism 9. Data for both crude venoms were normalised as above, then plotted as 95% confidence band curves using GraphPad Prism 9. IC50 (MTT) values were calculated using the ‘[Inhibitor] vs. normalized response -- Variable slope’ -- Variable slope’ function.

### SDS-PAGE gel electrophoresis

To visualise which proteins were present in each venom, reducing sodium dodecyl sulphate-polyacrylamide gel electrophoresis (SDS-PAGE) was performed using neat venom fractions. Briefly, 10 µL fraction in 10 µL reducing SDS-PAGE buffer (6X stock solution with 30% β-mercaptoethanol) was denatured at 100 °C for five minutes. Following this, 10 µL sample or protein marker (Thermo Scientific, 26616) per lane were loaded onto BioRad Mini-PROTEAN TGX precast (10 well, 4-20%) gels. Gels were run at 200 V for 30 minutes, before transfer into Coomassie blue solution (45% methanol, 45% distilled H2O, 10% acetic acid glacial, 0.25 g Coomassie blue to 100 mL) and left for one hour at room temperature on a shaker. These were rinsed with destain (45% methanol, 45% distilled H2O, 10% acetic acid glacial) and left in fresh destain overnight at room temperature on a shaker, before imaging on a GelDoc Go Imaging System (BioRad).

### Quantifying snake venom metalloproteinase activity

To quantify the SVMP activity of the various venom fractions, a previously described SVMP activity assay was performed [6,50]. Fractions of *E. romani* and *B. arietans* venom were used in duplicate, with *E. romani* fractions used at the concentration resulting from chromatographic separation, while *B. arietans* fractions were added at a 1:9 dilution (in PBS, pH 7.4) due to neat venom fractions resulting in signals too high for accurate quantification. Fifteen microlitres of each sample were added to a clear-sided, clear-, flat-bottomed 384-well plate, alongside a DMSO negative control. The ES010 SVMP substrate stock (6.2 mM in DMSO; R&D Systems) was diluted to 9.1 µM in Tris-HCl buffer (150 mM NaCl, 50 mM Tris-HCl; pH 7.5), before 75 µL was added to each well of the plate using a Viaflo 384 (Integra-biosciences), leading to a final concentration of 7.5uM once added to the wells. The plate was placed immediately into a CLARIOstar Plus (BMG Labtech) and read for ten minutes (Ex320 nm, Em420) using CLARIOstar Control software. The assay was conducted at room temperature. Generated %-RFU values for each fraction were normalized to the values obtained from crude venom. Data were plotted as bar graphs using GraphPad Prism 9.

## Statistical analysis

All data from cell-based assays are presented as mean average ± standard deviation of at least three independent experimental replicates. For cell experiments, ‘n’ is defined as an independent experiment completed at a separate time from other ‘n’s within that group of experiments; all drug and/or venom treatments within an ‘n’ were completed in triplicate wells and the mean taken as the final value for that one trial. All data from SVMP-substrate assays are presented as mean average ± standard deviation of two independent experimental replicates. Unpaired t-tests were performed for dual comparisons, one-way analysis of variances (ANOVAs) were performed for multiple comparisons with one independent variable followed by Dunnett’s multiple comparisons tests, as recommended by GraphPad Prism. A difference was considered statistically significant where *P* ≤ 0.05.

## Supplementary Materials

The following supporting information is available in the supplementary materials, Figure S1: Gel filtration chromatography of whole *E. romani* venom; Figure S2: Gel filtration chromatography of whole *B. arietans* venom.

## Author Contributions

Conceptualisation, KEB and NRC; Methodology, all authors; Formal analysis, KEB, AW, and MCW; Investigation, KEB, AW, and MCW; Data curation, KEB and MCW; Writing—original draft preparation, KEB, MCW, and NRC; Writing—review and editing, all authors; Visualisation, KEB and MCW; Supervision, NRC; Project administration, KEB and NRC; Funding acquisition, NRC.

## Data Availability Statement

Data are available at EMBL-EBI BioStudies (DOI:10.6019/S-BSST1746).

## Acknowledgments

We thank Paul Rowley and Edouard Crittenden for maintenance of snakes and provision of venom, and Dr Steven Hall for advice regarding the EDTA safety-testing in the cell model.

## References

1. Gutiérrez, J.M.; Calvete, J.J.; Habib, A.G.; Harrison, R.A.; Williams, D.J.; Warrell, D.A. Snakebite Envenoming. Nat. Rev. Dis. Primer 2017, 3, 17063, doi:10.1038/nrdp.2017.63.

2. World Health Organization (WHO) Snakebite Envenoming: A Strategy for Prevention and Control Available online: https://www.who.int/publications-detail/9789241515641 (accessed on 13 November 2024).

3. Ralph, R.; Sharma, S.K.; Faiz, M.A.; Ribeiro, I.; Rijal, S.; Chappuis, F.; Kuch, U. The Timing Is Right to End Snakebite Deaths in South Asia. BMJ 2019, 364, doi:10.1136/bmj.k5317.

4. Laustsen, A.H.; Johansen, K.H.; Engmark, M.; Andersen, M.R. Recombinant Snakebite Antivenoms: A Cost-Competitive Solution to a Neglected Tropical Disease? PLoS Negl. Trop. Dis. 2017, 11, e0005361, doi:10.1371/journal.pntd.0005361.

5. Ahmadi, S.; Pucca, M.B.; Jürgensen, J.A.; Janke, R.; Ledsgaard, L.; Schoof, E.M.; Sørensen, C.V.; Çalışkan, F.; Laustsen, A.H. An *in Vitro* Methodology for Discovering Broadly-Neutralizing Monoclonal Antibodies. Sci. Rep. 2020, 10, 10765, doi:10.1038/s41598-020-67654-7.

6. Albulescu, L.O.; Hale, M.S.; Ainsworth, S.; Alsolaiss, J.; Crittenden, E.; Calvete, J.J.; Evans, C.; Wilkinson, M.C.; Harrison, R.A.; Kool, J.;, et al. Preclinical Validation of a Repurposed Metal Chelator as an Early-Intervention Therapeutic for Hemotoxic Snakebite. Sci. Transl. Med. 2020, 12, eaay8314, doi:10.1126/scitranslmed.aay8314.

7. Lewin, M.R.; Carter, R.W.; Matteo, I.A.; Samuel, S.P.; Rao, S.; Fry, B.G.; Bickler, P.E. Varespladib in the Treatment of Snakebite Envenoming: Development History and Preclinical Evidence Supporting Advancement to Clinical Trials in Patients Bitten by Venomous Snakes. Toxins 2022, 14, doi:10.3390/toxins14110783.

8. Lauridsen, L.P.; Laustsen, A.H.; Lomonte, B.; Gutiérrez, J.M. Toxicovenomics and Antivenom Profiling of the Eastern Green Mamba Snake (*Dendroaspis Angusticeps*). J. Proteomics 2016, 136, 248–261, doi:10.1016/j.jprot.2016.02.003.

9. Ferraz, C.R.; Arrahman, A.; Xie, C.; Casewell, N.R.; Lewis, R.J.; Kool, J.; Cardoso, F.C. Multifunctional Toxins in Snake Venoms and Therapeutic Implications: From Pain to Hemorrhage and Necrosis. Front. Ecol. Evol. 2019, 7, doi:10.3389/fevo.2019.00218.

10. Liu, C.C.; Chou, Y.S.; Chen, C.Y.; Liu, K.L.; Huang, G.J.; Yu, J.S.; Wu, C.J.; Liaw, G.W.; Hsieh, C.H.; Chen, C. Pathogenesis of Local Necrosis Induced by *Naja Atra* Venom: Assessment of the Neutralization Ability of Taiwanese Freeze-Dried Neurotoxic Antivenom in Animal Models. PLoS Negl. Trop. Dis. 2020, 14, e0008054– e0008054, doi:10.1371/journal.pntd.0008054.

11. Chippaux, J.P. Estimate of the Burden of Snakebites in Sub-Saharan Africa: A Meta-Analytic Approach. Toxicon 2011, 57, 586–599, doi:10.1016/j.toxicon.2010.12.022.

12. World Health Organization (WHO) Guidelines for the Prevention and Clinical Management of Snakebite in Africa [Online: Accessed 11 November 2024]. 2010.

13. Megale, A.A.A.; Portaro, F.C.; Da Silva, W.D. *Bitis Arietans* Snake Venom Induces an Inflammatory Response Which Is Partially Dependent on Lipid Mediators. Toxins 2020, 12, 594, doi:10.3390/toxins12090594.

14. Pook, C.E.; Joger, U.; Stümpel, N.; Wüster, W. When Continents Collide: Phylogeny, Historical Biogeography and Systematics of the Medically Important Viper Genus *Echis* (Squamata: Serpentes: Viperidae). Mol. Phylogenet. Evol. 2009, 53, 792–807, doi:10.1016/j.ympev.2009.08.002.

15. Trape, JF. Partition d’Echis Ocellatus Stemmler, 1970 (Squamata, Viperidae), Avec La Description d’une Espèce Nouvelle. Bull. Société Herpétologique Fr. 2018, 167, 13–34.

16. Habib, A.G. Venomous Snakes and Snake Envenomation in Nigeria. In Clinical Toxinology in Asia Pacific and Africa; Gopalakrishnakone, P., Faiz, A., Fernando, R., Gnanathasan, C.A., Habib, A.G., Yang, C., Eds.; Toxinology; Springer: Dordrecht, 2015; Vol. 2, pp. 275–298.

17. Habib, A.G. Public Health Aspects of Snakebite Care in West Africa: Perspectives from Nigeria. J. Venom. Anim. Toxins Trop. Dis. 2013, 19, doi:10.1186/1678-9199-19-27.

18. Wagstaff, S.C.; Sanz, L.; Juarez, P.; Harrison, R.A.; Calvete, J.J. Combined Snake Venomics and Venom Gland Transcriptomic Analysis of the Ocellated Carpet Viper,*Echis Ocellatus*. J. Proteomics 2009, 71, 609–623, doi:10.1016/j.jprot.2008.10.003.

19. Casewell, N.R.; Wagstaff, S.C.; Wüster, W.; Cook, D.A.N.; Bolton, F.M.S.; King, S.I.; Pla, D. Medically Important Differences in Snake Venom Composition Are Dictated by Distinct Postgenomic Mechanisms. PNAS 2014, 111, 9205–9210, doi:10.1073/pnas.1405484111.

20. Tasoulis, T.; Isbister, G.K. A Review and Database of Snake Venom Proteomes. Toxins 2017, 9, 290, doi:10.3390/toxins9090290.

21. Dingwoke, E.J.; Adamude, F.A.; Mohamed, G.; Klein, A.; Salihu, A.; Abubakar, M.S.; Sallau, A.B. Venom Proteomic Analysis of Medically Important Nigerian Viper *Echis Ocellatus* and *Bitis Arietans* Snake Species. Biochem. Biophys. Rep. 2021, 28, doi:10.1016/j.bbrep.2021.101164.

22. Dawson, C.A.; Bartlett, K.E.; Wilkinson, M.C.; Ainsworth, S.; Albulescu, L.O.; Kazandjian, T.D.; Hall, S.R.; Westhorpe, A.P.; Clare, R.H.; Wagstaff, S.C.;, et al. Intraspecific Venom Variation in the Medically Important Puff Adder (*Bitis Arietans*): Comparative Venom Gland Transcriptomics, in Vitro Venom Activity and Immunological Recognition by Antivenom. PLoS Negl. Trop. Dis. 2024, doi:10.1371/journal.pntd.0012570.

23. Olaoba, O.T.; dos Santos, P.K.; Selistre-de-Araujo, H.S.; de Souza, D.H.F. Snake Venom Metalloproteinases (SVMPs): A Structure-Function Update. Toxicon X 2020, 100052, doi:10.1016/j.toxcx.2020.100052.

24. Hiu, J.J.; Yap, M.K.K. The Myth of Cobra Venom Cytotoxin: More than Just Direct Cytolytic Actions. Toxicon X 2022, 14, doi:10.1016/j.toxcx.2022.100123.

25. Escalante, T.; Rucavado, A.; Pinto, A.F.M.; Terra, R.M.S.; Gutiérrez, J.M.; Fox, J.W. Wound Exudate as a Proteomic Window to Reveal Different Mechanisms of Tissue Damage by Snake Venom Toxins. J. Proteome Res. 2009, 8, 5120– 5131, doi:10.1021/pr900489m.

26. Gutiérrez, J.M.; Escalante, T.; Rucavado, A.; Herrera, C.; Fox, J.W. A Comprehensive View of the Structural and Functional Alterations of Extracellular Matrix by Snake Venom Metalloproteinases (SVMPs): Novel Perspectives on the Pathophysiology of Envenoming. Toxins 2016, 8, 304, doi:10.3390/toxins8100304.

27. Slagboom, J.; Kool, J.; Harrison, R.A.; Casewell, N.R. Haemotoxic Snake Venoms: Their Functional Activity, Impact on Snakebite Victims and Pharmaceutical Promise. Br. J. Haematol. 2017, 177, 947–959, doi:10.1111/bjh.14591.

28. Fox, J.W.; Serrano, S.M.T. Insights into and Speculations about Snake Venom Metalloproteinase (SVMP) Synthesis, Folding and Disulfide Bond Formation and Their Contribution to Venom Complexity. FEBS J. 2008, 275, 3016–30 30, doi:10.1111/j.1742-4658.2008.06466.x.

29. Freitas-de-Sousa, L.A.; Colombini, M.; Lopes-Ferreira, M.; Serrano, S.M.T.; Moura-da-Silva, A.M. Insights into the Mechanisms Involved in Strong Hemorrhage and Dermonecrosis Induced by Atroxlysin-Ia, a PI-Class Snake Venom Metalloproteinase. Toxins 2017, 9, 239, doi:10.3390/toxins9080239.

30. Moretto Del-Rei, T.H.; Sousa, L.F.; Rocha, M.M.T.; Freitas-de-Sousa, L.A.; Travaglia-Cardoso, S.R.; Grego, K.; Sant’Anna, S.S.; Chalkidis, H.M.; Moura-da-Silva, A.M. Functional Variability of *Bothrops Atrox* Venoms from Three Distinct Areas across the Brazilian Amazon and Consequences for Human Envenomings. Toxicon 2019, 164, 61–70, doi:10.1016/j.toxicon.2019.04.001.

31. Khan, S.A.; Ilies, M.A. The Phospholipase A2 Superfamily: Structure, Isozymes, Catalysis, Physiologic and Pathologic Roles. Int. J. Mol. Sci. 2023, 24, 1353, doi:10.3390/ijms24021353.

32. Lomonte, B.; Angulo, Y.; Rufini, S.; Cho, W.; Giglio, J.R.; Ohno, M.; Daniele, J.J.; Geoghegan, P.; Gutiérrez, J.M. Comparative Study of the Cytolytic Activity of Myotoxic Phospholipases A2 on Mouse Endothelial (tEnd) and Skeletal Muscle (C2C12) Cells *in Vitro*. Toxicon 1999, 37, 145–158, doi:10.1016/S0041-0101(98)00171-8.

33. Gasanov, S.E.; Dagda, R.K.; Rael, E.D. Snake Venom Cytotoxins, Phospholipase A2s, and Zn2+-Dependent Metalloproteinases: Mechanisms of Action and Pharmacological Relevance. J. Clin. Toxicol. 2014, 4, 1000181– 1000181, doi:10.4172/2161-0495.1000181.

34. Araya, C.; Lomonte, B. Antitumor Effects of Cationic Synthetic Peptides Derived from Lys49 Phospholipase A2 Homologues of Snake Venoms. Cell Biol. Int. 2007, 31, 263–268, doi:10.1016/j.cellbi.2006.11.007.

35. Bryan-Quiros, W.; Fernandez, J.; Gutierrez, J.M.; Lewin, M.R.; Lomonte, B. Neutralizing Properties of LY315920 toward Snake Venom Group I and II Myotoxic Phospholipases A2. Toxicon 2019, 157, 1–7, doi:10.1016/j.toxicon.2018.11.292.

36. Ghazaryan, N.A.; Ghulikyan, L.; Kishmiryan, A.; Andreeva, T.V.; Utkin, Y.N.; Tsetlin, V.I.; Lomonte, B.; Ayvazyan, N.M. Phospholipases A2 from Viperidae Snakes: Differences in Membranotropic Activity between Enzymatically Active Toxin and Its Inactive Isoforms. Biochim. Biophys. Acta 2015, 1848, 463–468, doi:10.1016/j.bbamem.2014.10.037.

37. Fernández, J.; Gutiérrez, J.M.; Angulo, Y.; Sanz, L.; Juárez, P.; Calvete, J.J.; Lomonte, B. Isolation of an Acidic Phospholipase A2 from the Venom of the Snake *Bothrops Asper* of Costa Rica: Biochemical and Toxicological Characterization. Biochimie 2010, 92, 273–283, doi:10.1016/j.biochi.2009.12.006.

38. Kang, T.S.; Georgieva, D.; Genov, N.; Murakami, M.T.; Sinha, M.; Kumar, R.P.; Kaur, P.; Kumar, S.; Dey, S.; Sharma, S.;, et al. Enzymatic Toxins from Snake Venom: Structural Characterization and Mechanism of Catalysis. FEBS J. 2011, 278, 4544–4576, doi:10.1111/j.1742-4658.2011.08115.x.

39. Costal-Oliveira, F.; Stransky, S.; Guerra-Duarte, C.; Naves de Souza, D.L.; Vivas-Ruiz, D.E.; Yarlequé, A.; Sanchez, E.F.; Chávez-Olórtegui, C.; Braga, V.M.M. L-Amino Acid Oxidase from *Bothrops Atrox* Snake Venom Triggers Autophagy, Apoptosis and Necrosis in Normal Human Keratinocytes. Sci. Rep. 2019, 9, 1–14, doi:10.1038/s41598-018-37435-4.

40. Abdelkafi-Koubaa, Z.; Elbini-Dhouib, I.; Souid, S.; Jebali, J.; Doghri, R.; Srairi-Abid, N.; Essafi-Benkhadir, K.; Micheau, O.; Marrakchi, N. Pharmacological Investigation of CC-LAAO, an L-Amino Acid Oxidase from *Cerastes Cerastes* Snake Venom. Toxins 2021, 13, 904, doi:10.3390/toxins13120904.

41. Bartlett, K.E.; Hall, S.R.; Rasmussen, S.A.; Crittenden, E.; Dawson, C.A.; Albulescu, L.O.; Laprade, W.; Harrison, R.A.; Saviola, A.J.; Modahl, C.M.;, et al. Dermonecrosis Caused by Spitting Cobra Snakebite Is the Result of Toxin Potentiation and Is Prevented by the Repurposed Drug Varespladib. PNAS 2024, 121, e2315597121, doi:10.1073/pnas.2315597121.

42. Hall, S.R.; Rasmussen, S.A.; Crittenden, E.; Dawson, C.A.; Bartlett, K.E.; Westhorpe, A.P.; Albulescu, L.; Kool, J.; Gutiérrez, J.M.; Casewell, N.R. Repurposed Drugs and Their Combinations Prevent Morbidity-Inducing Dermonecrosis Caused by Diverse Cytotoxic Snake Venoms. Nat. Commun. 2023, 14, 7812, doi:10.1038/s41467-023-43510-w.

43. Mosmann, T. Rapid Colorimetric Assay for Cellular Growth and Survival: Application to Proliferation and Cytotoxicity Assays. J. Immunol. Methods 1983, 65, 55–63, doi:10.1016/0022-1759(83)90303-4.

44. Fotakis, G.; Timbrell, J.A. *In Vitro* Cytotoxicity Assays: Comparison of LDH, Neutral Red, MTT and Protein Assay in Hepatoma Cell Lines Following Exposure to Cadmium Chloride. Toxicol. Lett. 2006, 160, 171–177, doi:10.1016/j.toxlet.2005.07.001.

45 Wilkinson, M.C.; Modahl, C.; Saviola, A.; Albulesco, L.O.; Tianyi, F.L.; Casewell, N.R. Isolation and Characterisation of Serine Proteases and Metalloproteases from the Venom of African Puff Adders [Preprint]. BioRxiv 2024, doi:10.1101/2024.05.31.596867.

46. Ainsworth, S.; Slagboom, J.; Alomran, N.; Pla, D.; Alhamdi, Y.; King, S.I.; Bolton, F.M.S. The Paraspecific Neutralisation of Snake Venom Induced Coagulopathy by Antivenoms. *Commun*. Biol. 2018, 1, doi:doi:10.1038/s42003-018-0039-1.

47. Borkow, G.; Gutiérrez, J.M.; Ovadia, M. Inhibition of the Hemorrhagic Activity of *Bothrops Asper* Venom by a Novel Neutralizing Mixture. Toxicon 1997, 35, 865–877, doi:10.1016/S0041-0101(96)00193-6.

48. Gutiérrez, J.M.; Chaves, F.; Rucavado, A.; León, G.; Franceschi, A.; Ovadia, M.; Escalante, T.; Cury, Y. Inhibition of Local Hemorrhage and Dermonecrosis Induced by *Bothrops Asper* Snake Venom: Effectiveness of Early in Situ Administration of the Peptidomimetic Metalloproteinase Inhibitor Batimastat and the Chelating Agent CaNa2EDTA. Am. J. Trop. Med. Hyg. 2000, 63, 313–319, doi:10.4269/ajtmh.2000.63.313.

49. Howes, J.M.; Theakston, R.D.G.; Laing, G.D. Neutralization of the Haemorrhagic Activities of Viperine Snake Venoms and Venom Metalloproteinases Using Synthetic Peptide Inhibitors and Chelators. Toxicon 2007, 49, 734– 739, doi:10.1016/j.toxicon.2006.11.020.

50. Menzies, S.K.; Clare, R.H.; Xie, C.; Westhorpe, A.; Hall, S.R.; Edge, R.J.; Alsolaiss, J.; Crittenden, E.; Marriott, A.E.; Harrison, R.A.;, et al. *In Vitro* and *in Vivo* Preclinical Venom Inhibition Assays Identify Metalloproteinase Inhibiting Drugs as Potential Future Treatments for Snakebite Envenoming by *Dispholidus Typus*. Toxicon X 2022, 14, 100118, doi:10.1016/j.toxcx.2022.100118.

51. Currier, R.B.; Harrison, R.A.; Rowley, P.D.; Laing, G.D.; Wagstaff, S.C. Intra-Specific Variation in Venom of the African Puff Adder (*Bitis Arietans*): Differential Expression and Activity of Snake Venom Metalloproteinases (SVMPs). Toxicon Off. J. Int. Soc. Toxinology 2010, 55, 864–873, doi:10.1016/j.toxicon.2009.12.009.

52. Warrell, D.A.; Williams, D.J. Clinical Aspects of Snakebite Envenoming and Its Treatment in Low-Resource Settings. The Lancet 2023, doi:10.1016/S0140-6736(23)00002-8.

53. Gutiérrez, J.M.; Casewell, N.R.; Laustsen, A.H. Progress and Challenges in the Field of Snakebite Envenoming Therapeutics. Annu. Rev. Pharmacol. Toxicol. 2024, doi:10.1146/annurev-pharmtox-022024-033544.

54. Warrell, D.A.; Davidson, N.M.; Greenwood, B.M.; Ormerod, L.D.; Pope, H.M.; Watkins, B.J.; Prentice, C.R.M. Poisoning by Bites of the Saw-Scaled or Carpet Viper (*Echis Carinatus*) in Nigeria. *QJM Int*. J. Med. 1977, 46, 33–62, doi:10.1093/oxfordjournals.qjmed.a067493.

55. Fernandez, S.; Hodgson, W.; Chaisakul, J.; Kornhauser, R.; Konstantakopoulos, N.; Smith, A.I.; Kuruppu, S. *In Vitro* Toxic Effects of Puff Adder (*Bitis Arietans*) Venom, and Their Neutralization by Antivenom. Toxins 2014, 6, 1586–1597, doi:10.3390/toxins6051586.

56. Clare, R.H.; Hall, S.R.; Patel, R.N.; Casewell, N.R. Small Molecule Drug Discovery for Neglected Tropical Snakebite. Trends Pharmacol. Sci. 2021, 42, 340–353, doi:10.1016/j.tips.2021.02.005.

57. Sánchez, A.; Coto, J.; Segura, Á.; Vargas, M.; Solano, G.; Herrera, M.; Villalta, M.; Estrada, R.; Gutiérrez, J.M.; León,

58. G. Effect of Geographical Variation of *Echis Ocellatus*, *Naja Nigricollis* and *Bitis Arietans* Venoms on Their Neutralization by Homologous and Heterologous Antivenoms. Toxicon 2015, 108, 80–83, doi:10.1016/j.toxicon.2015.10.001.

58. Bhatia, S.; Vasudevan, K. Comparative Proteomics of Geographically Distinct Saw-Scaled Viper (*Echis Carinatus*) Venoms from India. Toxicon X 2020, 7, 100048, doi:10.1016/j.toxcx.2020.100048.

59. Senji, L.R.R.; Attarde, S.; Khochare, S.; Suranse, V.; Martin, G.; Casewell, N.R.; Whitaker, R.; Sunagar, K. Biogeographical Venom Variation in the Indian Spectacled Cobra (*Naja Naja*) Underscores the Pressing Need for Pan-India Efficacious Snakebite Therapy. PLoS Negl. Trop. Dis. 2021, 15, e0009150, doi:10.1371/journal.pntd.0009150.

60. Mora-Obando, D.; Salazar-Valenzuela, D.; Pla, D.; Lomonte, B.; Guerrero-Vargas, J.A.; Ayerbe, S.; Gibbs, H.L.; Calvete, J.J. Venom Variation in *Bothrops Asper* Lineages from North-Western South America. J. Proteomics 2020, 229, 103945, doi:10.1016/j.jprot.2020.103945.

61. Silva-Júnior, L.N.; Abreu, L.D.S.; Rodrigues, C.F.B.; Galizio, N.D.C.; Aguiar, W.D.S.; Serino-Silva, C.; Santos, V.S.D.; Costa, I.A.; Oliveira, L.V.F.; Sant’Anna, S.S.;, et al. Geographic Variation of Individual Venom Profile of *Crotalus Durissus* Snakes. J. Venom. Anim. Toxins Trop. Dis. 2020, 26, e20200016, doi:10.1590/1678-9199-jvatitd-2020-0016.

62. Smith, C.F.; Nikolakis, Z.L.; Ivey, K.; Perry, B.W.; Schield, D.R.; Balchan, N.R.; Parker, J.; Hansen, K.C.; Saviola, A.J.; Castoe, T.A.;, et al. Snakes on a Plain: Biotic and Abiotic Factors Determine Venom Compositional Variation in a Wide-Ranging Generalist Rattlesnake. BMC Biol. 2023, 21, 136, doi:10.1186/s12915-023-01626-x.

63. Tianyi, F.L.; Ngari, C.; Wilkinson, M.C.; Parkurito, S.; Chebet, E.; Mumo, E.; Trelfa, A.; Otundo, D.; Crittenden, E.; Kephah, G.M.;, et al. Clinical Features of Puff Adder Envenoming: Case Series of *Bitis Arietans* Snakebites in Kenya and a Review of the Literature [Preprint]. MedRxiv 2024, doi:10.1101/2024.05.31.24308288.

64. Zancolli, G.; Baker, T.; Barlow, A.; Bradley, R.; Calvete, J.; Carter, K.; De Jager, K.; Owens, J.; Price, J.; Sanz, L.;, et al. Is Hybridization a Source of Adaptive Venom Variation in Rattlesnakes? A Test, Using a *Crotalus Scutulatus* × *Viridis* Hybrid Zone in Southwestern New Mexico. Toxins 2016, 8, 188, doi:10.3390/toxins8060188.

65. Casewell, N.R.; Jackson, T.N.W.; Laustsen, A.H.; Sunagar, K. Causes and Consequences of Snake Venom Variation. Trends Pharmacol. Sci. 2020, 41, 570–581, doi:10.1016/j.tips.2020.05.006.

66. Wilson, V.G. Growth and Differentiation of HaCaT Keratinocytes. Methods Mol. Biol. 2014, 1195, doi:10.1007/7651_2013_42.

67. Colombo, I.; Sangiovanni, E.; Maggio, R.; Mattozzi, C.; Zava, S.; Corbett, Y.; Fumagalli, M.; Carlino, C.; Corsetto, P.A.; Scaccabarozzi, D.;, et al. HaCaT Cells as a Reliable *in Vitro* Differentiation Model to Dissect the Inflammatory/Repair Response of Human Keratinocytes. Mediators Inflamm. 2017, 2017, doi:10.1155/2017/7435621.

