## Supplementary material for "Snake venom metalloproteinases are predominantly responsible for the cytotoxic effects of certain African viper venoms": Figure S1

**Keirah E. Bartlett [1,#], Adam Westhorpe [1], Mark C. Wilkinson [1] and Nicholas R. Caswell [1,*]**

1 Centre for Snakebite Research & Interventions, Department of Tropical Disease Biology, Liverpool School of Tropical Medicine, Liverpool L3 5QA, United Kingdom
### Current address: School of Pharmacy, University of Reading, Reading RG6 6AU, United Kingdom

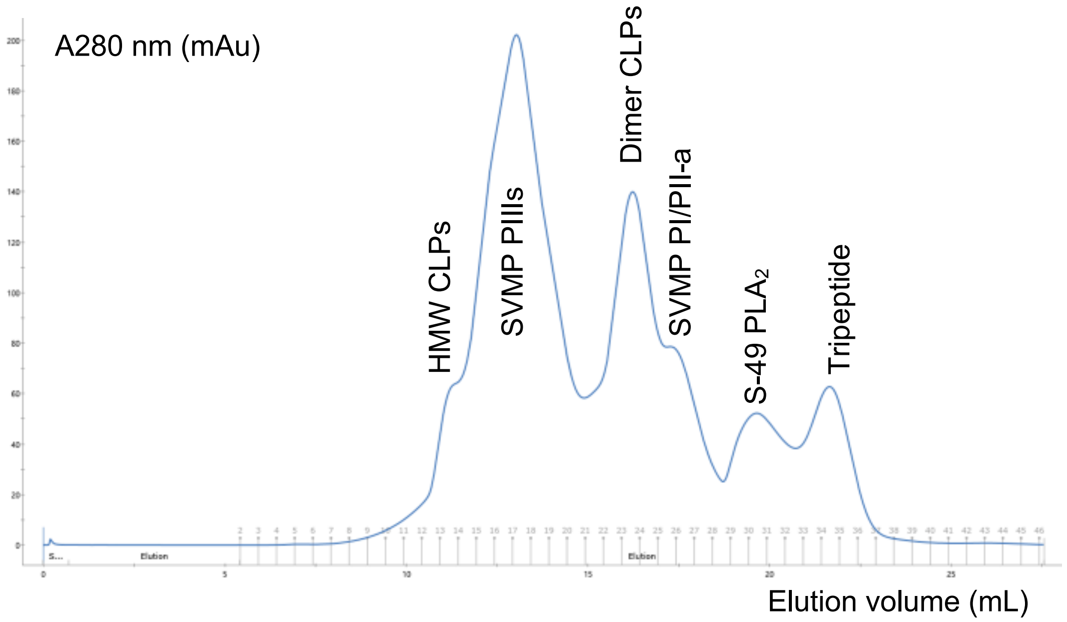


**Figure S1. Gel filtration chromatography of whole *E. romani* venom.** A 24 mL Superdex 200 column was used for the separation which was pre-equilibrated in PBS [25 mM sodium phosphate, 0.15 M NaCl, pH 7.2]. A 0.5 mL aliquot of *E. romani* venom at a concentration of 10 mg/mL in PBS was injected onto the column and the separation was carried out in PBS at a flow-rate of 0.3 mL/min. Elution was monitored at 280 nm and 0.5 mL fractions were collected. [HMW = high molecular weight].

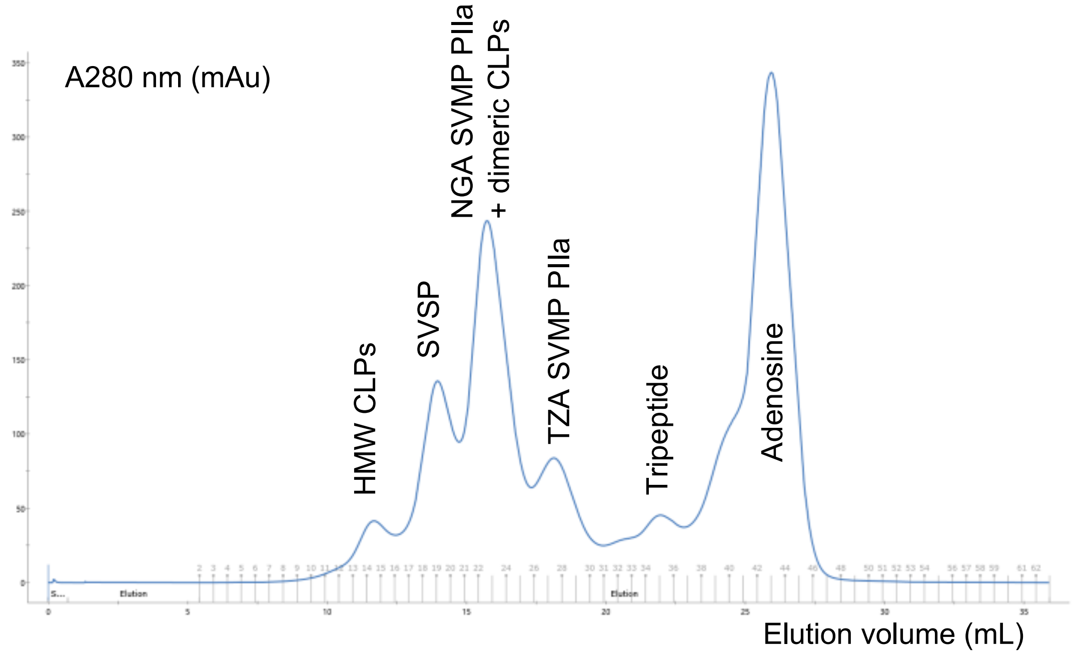


**Figure S2. Gel filtration chromatography of whole *B. arietans* venom.** A 24 mL Superdex 200 column was used for the separation which was pre-equilibrated in PBS [25 mM sodium phosphate, 0.15 M NaCl, pH 7.2]. A 0.5 mL aliquot of *B. arietans* venom at a concentration of 10 mg/mL in PBS was injected onto the column and the separation was carried out in PBS at a flow-rate of 0.3 mL/min. Elution was monitored at 280 nm and 0.5 mL fractions were collected. [HMW = high molecular weight; CLP = C-type lectin like protein; SVSP = serine protein, NGA = Nigerian; TZA = Tanzanian]
